# Intein-based thermoregulated meganucleases for biocontainment of genetic material

**DOI:** 10.1101/2023.08.14.553307

**Authors:** Gary W. Foo, Christopher D. Leichthammer, Ibrahim M. Saita, Nicholas D. Lukas, Izabela Z. Batko, David E. Heinrichs, David R. Edgell

## Abstract

Limiting the spread of synthetic genetic information outside of the intended use is essential for applications where biocontainment is critical. In particular, biocontainment of engineered probiotics and plasmids that are excreted from the mammalian gastrointestinal tract is needed to prevent escape and acquisition of genetic material that could confer a selective advantage to microbial communities. Here, we built a simple and lightweight biocontainment system that post-translationally activates a site-specific DNA endonuclease to degrade DNA at 18°C and not at higher temperatures. We constructed an orthogonal set of temperature sensitive-meganucleases, or TSMs, by inserting the yeast VMA1 L212P temperature-sensitive intein into the coding regions of LAGLIDADG homing endonucleases. We showed that the TSMs eliminated plasmids carrying the cognate TSM target site from laboratory strains of *Escherichia coli* at the permissive 18°C but not at higher restrictive temperatures. Plasmid elimination is dependent on both TSM endonuclease activity and intein splicing. We demonstrated that TSMs eliminated plasmids from the *E. coli* Nissle 1917 strain after passage through the mouse gut when fecal resuspensions were incubated at 18°C but not at 37°C. Collectively, our data demonstrates the potential of thermoregulated meganucleases as a means of restricting engineered plasmids and probiotics to the mammalian gut.

## Introduction

Biocontainment strategies to limit the spread of genetically modified microorganisms and associated genetic material are an essential component of synthetic biology research^1, 2^. In recent years, engineered bacterial probiotic strains have been designed to act as next-generation therapeutics to restore microbial homeostasis in dysbiotic conditions in the human microbiome^3–5^. For engineered strains that carry plasmids or other accessory elements, robust biocontainment systems are needed to prevent the escape and dissemination of recombinant genetic material before widespread adoption of engineered strains in human medicine, or other applications where biocontainment is critical. In particular, DNA released to the environment could be acquired by naturally competent bacteria or spread by mechanisms that promote lateral gene transfer, potentially conferring a selective advantage depending on the integrity and information carried on synthetic molecules^6–8^. Biocontainment systems must therefore be able to sufficiently degrade recombinant genetic material and limit acquisition by microbial communities^9–12^.

Current biocontainment strategies that rely on auxotrophic dependencies or self-killing systems that are activated by external signalling molecules show promise, but are optimized for laboratory environments and are not designed to limit environmental escape of genetic material after cell death. In contrast, kill switches comprised of site-specific or non-specific DNA endonucleases that degrade DNA upon activation are well suited to limit the spread of genetic material^10–12^. Previously, type II restriction enzymes such as EcoRI were used in endonuclease-based kill-switches and temperature regulated kill-switches based on the CRISPR/Cas9 system have also been developed^13, 14^. However, a single mutation in the sgRNA sequence can result in either loss of function or the creation of off-target sites^15,16^. This shortcoming could be overcome by multiplexing sgRNAs^17^, but with a concomitant increase in the complexity of the system, the number of potential targets for escape mutants, and the probability of off-target cleavage. Furthermore, many biocontainment systems are regulated by multi-layered genetic circuits controlled by exogenously added molecules, thus providing multiple points of potential mutational inactivation. Moreover, for applications in the mammalian GI tract, some of the required signalling molecules are potential carbon sources for the gut microbiome.

Here, we use a different biocontainment approach that is based on temperature as a trigger to post-translationally activate an endonuclease-based kill switch (Fig. 1). Our system is based on the insertion of the temperature-sensitive VMA1 L212P intein into the coding region of LAGLI-DADG homing endonucleases (we call these enzymes temperature-sensitive meganucleases, or TSMs)^18–23^. The VMA1 L212P intein splices at temperatures below 20°C^21^to activate TSMs for site-specific cleavage, eliminating plasmids carrying the cognate TSM target site from laboratory strains of *Escherichia coli*. TSMs also eliminated plasmids from the probiotic *E. coli* Nissle 1917 strain that had been passaged through the mouse gut when fecal resuspensions were incubated at 18°C but not at 37°C. Our data highlight the utility of a simple, post-translationally activated kill switch for temperature-regulated biocontainment.

**Figure 1.**
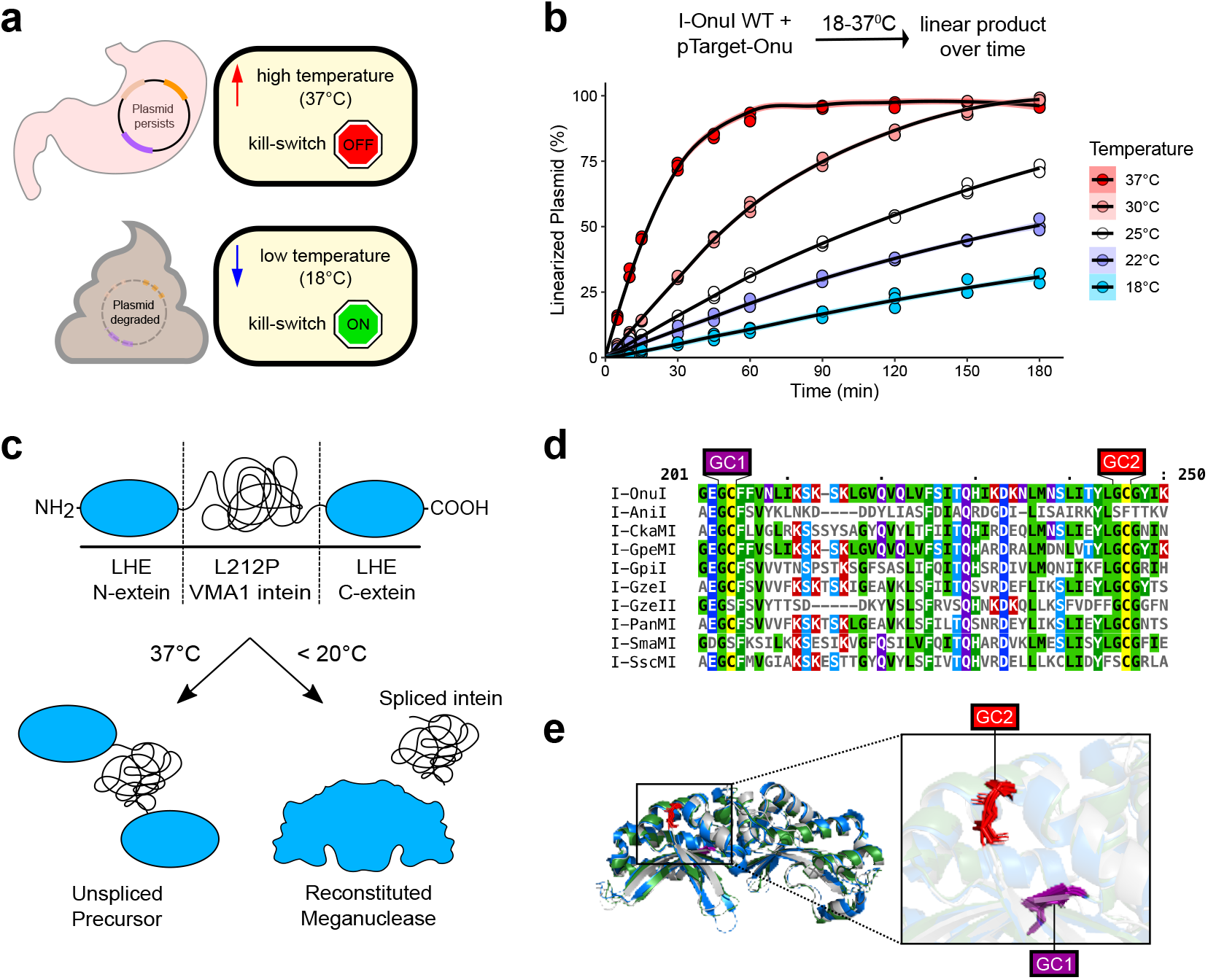
Strategy for the development of a thermosensitive endonuclease kill-switch. **(A)** Schematic of the thermosensitive kill-switch shown in the context of plasmid containment in the mammalian gut. **(B)** Linear product formation over time *in vitro* for wild-type I-OnuI incubated at different temperatures with pTarget-Onu. Each point represents an independent replicate (n=3). **(C)** Thermoregulation of LAGLIDADG homing endonucleases (LHEs) using the L212P VMA1 intein. (top) The VMA1/LHE open reading frame (not to scale) with the meaganuclease (LHE) N- and C-exteins and the VMA1 intein indicated. (bottom) Formation of an active meganuclease is dependent on low temperature to promote splicing of the VMA1 L212P intein, whereas higher temperatures inhibit splicing and formation of an active meganuclease. **(D)** Multiple sequence alignment of 10 LHEs with the two conserved glycine-cysteine sites for the insertion of the VMA1 intein labelled as GC1 and GC2 in purple and red, respectively. **(E)** Alignment of the crystal structures of I-OnuI, I-GpeMI, and I-PanMI. I-OnuI is shown in blue (PDB:6BDA), I-GpeMI in green (PDB:4YHX), and I-PanMI in silver (PDB:5ESP). GC1 and GC2 sites are highlighted in purple and red, respectively.

## Results

### Design of intein-based thermosensitive meganucleases

Our goal was to design a thermoregulated kill-switch activated at low permissive temperatures outside of the mammalian GI tract to promote elimination of recombinant plasmids, but inactive at higher restrictive temperatures while passaging through the GI tract (Fig. 1a). To this end, we selected LAGLIDADG homing endonucleases (LHEs, also called meganucleases) as candidate kill switches because of their small coding size, long 22-bp target sequences, tolerance to mutational inactivation, and structural conservation^24,25^. We first tested the cleavage activity of the well-characterized I-OnuI meganuclease^26^to determine if the enzyme was active at low temperatures (Fig. 1b, Fig. S1). Using a plasmid that contained the I-OnuI cognate target site (pTarget-Onu), we found I-OnuI was active over at least 5 temperatures spanning the 37°C to 18°C range but with a 4-fold reduction in activity at 18°C as compared to 37°C. This data suggests that I-OnuI would be appropriate for a kill switch if the activity could be regulated to low temperatures.

To create a thermoregulated version of I-OnuI, as well as other meganucleases, we re-purposed the *Saccharomyces cerevisiae* vacuolar ATPase (VMA1) L212P intein that splices at or below 18°C to control meganuclease activity (Fig. 1c)^20–23^. This post-translational activation of a kill switch differs from previous designs that rely on transcriptional control regulated by exogenously added signalling molecules. The VMA1 intein splices most efficiently when inserted downstream of a glycine (G) residue and upstream of a cysteine (C)^27–29^, and is enhanced when inserted in a loop region or proximal to the active site of the host protein^30^. We found two glycine-cysteine (GC) sites that are conserved amongst 10 aligned meganucleases, with at least one site being found in each meganuclease (Fig. 1d). All 10 meganucleases are functionally orthogonal, cleaving different cognate DNA substrates (Fig. S2). I-OnuI, I-GpeMI, and I-PanMI possessed both of the GC sites and aligning the crystal structures revealed that the GC sites are located in the C-terminal domain of each enzyme. The first GC site (GC1) present in a *β* -sheet that forms part of the I-OnuI protein-DNA interface while the second GC site (GC2) is present in an exposed alpha helix (Fig. 1e). We independently inserted the temperature sensitive L212P (TS) and wild-type (WT) VMA1 inteins into I-OnuI, I-GpeMI and I-PanMI. Hereafter, the meganuclease constructs with the L212P temperature-sensitive intein insertion are referred to as TSMs (thermosensitive meganucleases).

### Testing the thermosensitivity of TSMs

We next designed an *E. coli* two-plasmid assay to assess the thermosensitivity of meganucleases and TSMs. The assay included pEndo that expressed either the meganuclease or TSM from an arabinose-regulated promoter^31^and pTarget that contained the meganuclease target site. Active meganucleases/TSMs will cleave their cognate target sites on pTarget to promote elimination of pTarget by the RecBCD pathway^32^and subsequent loss of kanamycin resistance, with the TSMs only being active at low temperatures (Fig. 2a).

**Figure 2.**
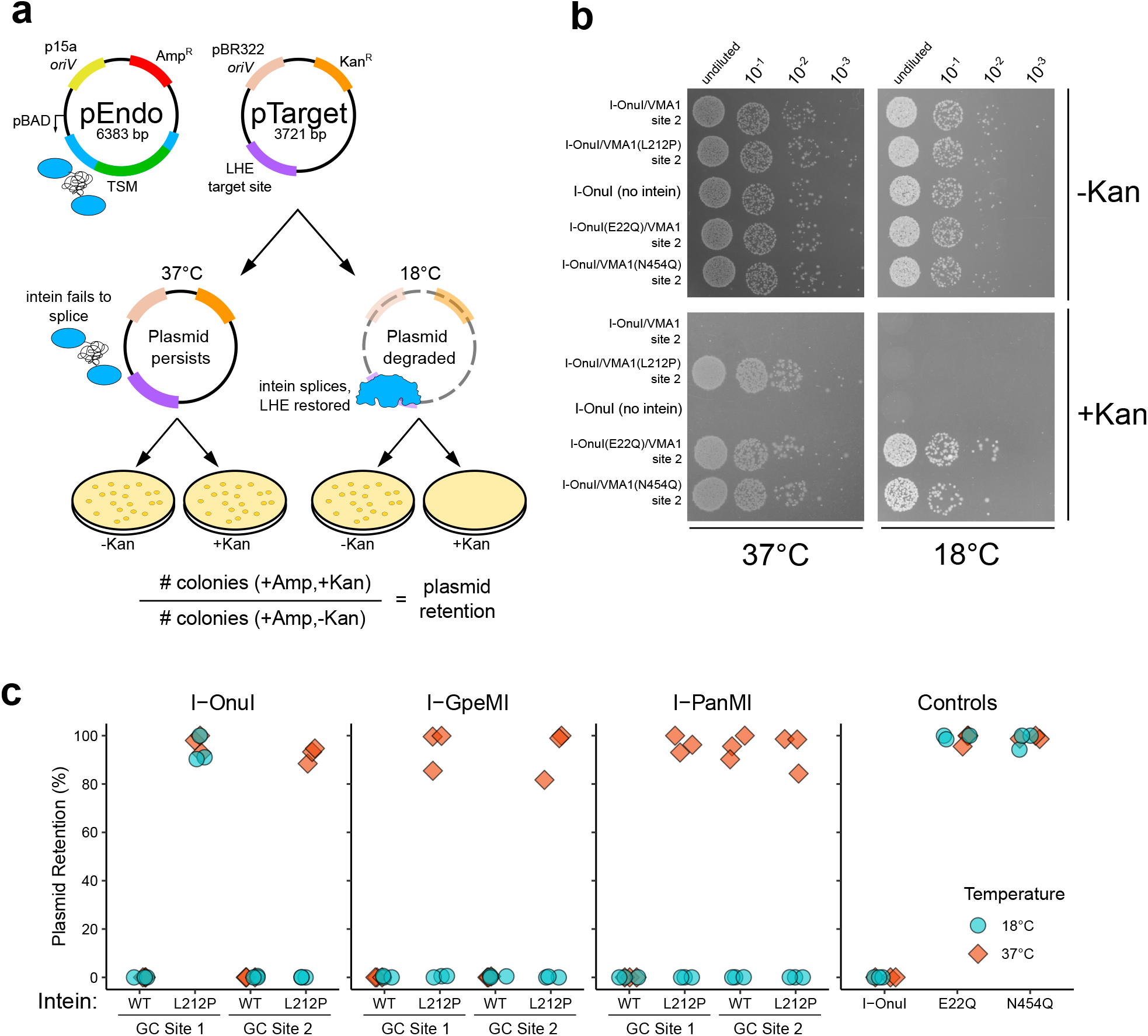
Thermoregulation of endonuclease activity using an intein-based approach. **(A)** Schematic of pEndo and pTarget used in the two-plasmid selection experiments. *oriV*, plasmid origin of replication; pBAD, arabinose-inducible promoter; TSM, temperature-sensitive meganuclease; Amp^R^, ampicillin resistance gene; Kan^R^, kanamycin resistance gene. The LHE target site is cleaved by the spliced TSM. **(B)** I-OnuI TSM cleavage activity in *E. coli* NEB 5-alpha on M9 minimal media incubated at either 37°C or 18°C. **(C)** Two-plasmid assay in *E. coli* NEB 5-alpha with TSMs developed from three LHEs. Plasmid retention was calculated as the ratio of colonies grown on M9 minimal media with ampicillin and kanamycin, over media with only ampicillin. Each data point represents individual replicates (n=3).

We first cloned the I-OnuI GC2 TSM into pEndo and assessed elimination of pTarget-Onu by spot plating over 1000-fold dilutions at 18°C and 37°C (Fig. 2b). We observed no growth at 18°C and robust growth at 37°C (Fig. 2b). We also determined pTarget-Onu retention by the ratio of colony counts on media containing or lacking kanamycin and observed no pTarget-Onu retention at 18°C as compared to near 100% retention at 37°C (Fig. 2c). These data are consistent with the L212P intein splicing at 18°C to produce a functional I-OnuI meganuclease that promotes elimination of pTarget and subsequent sensitivity to kanamycin. To confirm these observations, we performed parallel experiments with the WT VMA1 intein inserted in I-OnuI GC2, finding no growth at either 18°or 37°, indicating that splicing at both temperatures created a functional I-OnuI to eliminate pTarget (Fig. 2b and 2c). In contrast, the wild-type I-OnuI meganuclease with no intein insertion promoted pTarget-Onu elimination at either 18°C and 37°C (Fig. 2b and Fig. 2c). We also created endonuclease dead (E22Q) and intein-splicing dead (N454Q) variants in the I-OnuI GC2 construct. These variants promoted growth at either temperature, demonstrating that plasmid elimination is dependent on both endonuclease activity and intein splicing (Fig. 2b and 2c). Intein splicing did not create a toxic meganuclease at either temperature as we observed robust growth on plates lacking kanamycin.

To extend these findings to the second GC site in I-OnuI, and to other meganucleases, we created versions of pEndo carrying meganuclease variants as well as pTarget variants with the cognate target sites for each meganuclease. As shown in Fig. 2c, the insertion of the L212P VMA1 intein into the GC1 and GC2 sites of I-PanMI and I-GpeMI generated TSMs, whereas only the I-OnuI GC2 site generated a TSM. Interestingly, inserting the wild-type VMA1 intein into GC2 in I-PanMI generated a thermosensitive phenotype. As seen for the spot plating data (Fig. 2b), we observed near 100% plasmid retention for variants with substitutions that inactivated I-OnuI endonuclease activity (E22Q) and intein splicing (N454Q) (Fig. 2c). Orthogonality of the system was demonstrated by co-transforming pEndo programmed with the I-OnuI TSM and pTarget carrying the I-GpeMI site (pTarget-Gpe); pTarget-Gpe retention was *∼* 100% at 18°C, indicating lack of cleavage by I-OnuI (Fig. S3). Collectively, these experiments demonstrated that a set of orthogonal temperature-sensitive meganucleases (TSMs) were created by the insertion of the L212P VMA1 intein variant at splicing-permissive GC sites in I-OnuI, I-PanMI and I-GpeMI.

### Kinetics of TSM kill-switch activation

We next sought to characterize the I-OnuI GC2 TSM by determining the kinetics at the optimal restrictive temperature. We found that the I-OnuI TSM was fully functional at 18°C, partially functional at 20°C and exhibited a small-colony phenotype on solid media plates, and was inactive at higher temperatures (Fig. 3a). The small-colony phenotype was likely the result of incomplete pTarget-Onu cleavage, resulting in partial sensitivity to kanamycin. Further profiling was performed at 18°C.

**Figure 3.**
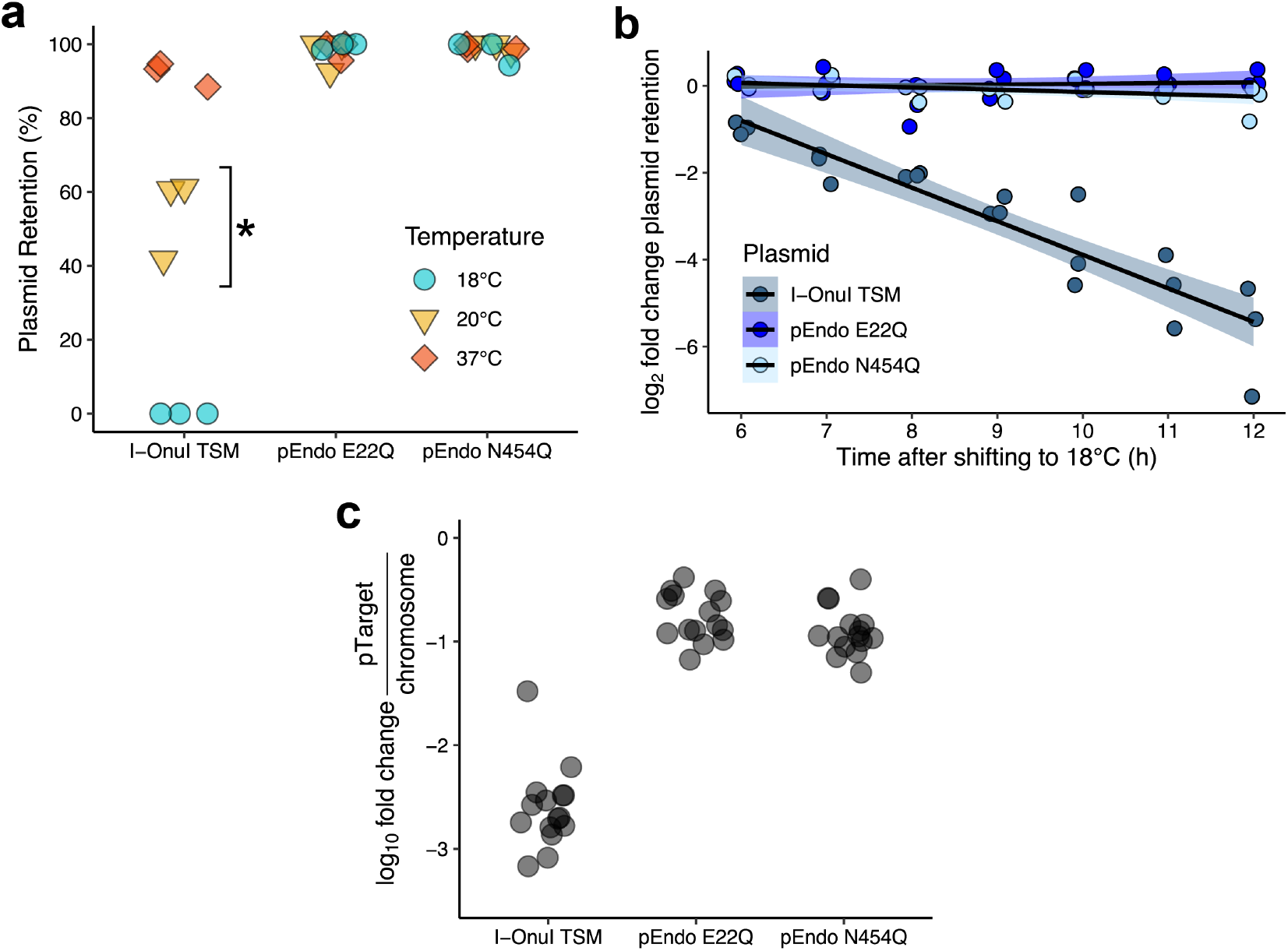
Enzyme kinetics and thermosensitivity of the I-OnuI TSM in *E. coli* Nissle 1917. **(A)** Two-plasmid assay with the I-OnuI TSM to determine the temperature at which plasmid cleavage is no longer observed. Each data point represents individual replicates (n=3). ^*^small colony morphology relative to the negative controls was observed on solid LB media incubated at 20°C. **(B)** *in vivo* cleavage of pTarget in liquid culture over time. Aliquots of the liquid culture were taken at the specified time points and plated on solid LB media, plasmid retention was calculated as previously described. Each data point represents individual replicates (n=3). **(C)** Change in the quantity of target plasmid relative to *E. coli* Nissle 1917 chromosome number after 12 hours of incubation at 18°C determined by quantitative PCR. Three biological and five technical replicates were performed for each sample, and each data point represents one of the replicates.

We next profiled the *in vivo* kinetics of kill-switch activation by diluting saturated cultures containing I-OnuI TSM and pTarget-Onu into fresh media lacking kanamycin and incubating the cultures at 18°C for up to 12 hours. At every hour, an aliquot was plated on solid media with or without kanamycin to calculate pTarget-Onu depletion. For these experiments, we used the probiotic strain *E. coli* Nissle 1917 (EcN) that has GRAS (generally recognized as safe) status instead of the NEB5*α* strain^33–36^. We also removed the pBAD regulatory cassette from pEndo, replacing it with the weak, constitutively active Anderson promoter (BBaJ23108) in anticipation of limiting complications of using sugar-related promoters for *in vivo* mouse experiments. With the I-OnuI TSM, minimal pTarget-Onu depletion was observed over the first six hours. In contrast, from 6-12 hours after shifting to 18°C, pTarget-Onu was eliminated at a linear rate (Fig. 3b). We observed no measurable pTarget-Onu elimination with cultures containing the pEndo N454Q intein splicing mutant or the I-OnuI E22Q endonuclease dead mutant (Fig. 3b). We estimate that by the end of the 12-hour incubation period, the I-OnuI TSM kill-switch had a 50-fold rate of pTarget-Onu depletion over both the negative controls.

To provide an independent measure of the I-OnuI TSM activity, we performed quantitative PCR on total DNA isolated after 12 hours at 18°C to determine the change in pTarget-Onu abundance relative to the *cspA* gene on the chromosome. The I-OnuI TSM reduced pTarget-Onu abundance *∼* 1000-fold *in vivo* as compared to EcN with the I-OnuI E22Q endonuclease dead and VMA1 N45Q splicing dead constructs (Fig. 3c). Taken together, these data show that I-OnuI TSM activity is highest at 18°C but that at least 6 hours incubation *in vivo* is required before clearance of pTarget-Onu can be observed.

### Multiplexing TSMs reduces the frequency of kill-switch escape mutants

An important characteristic of a biocontainment system is the rate of escape. In our system, escape would be characterized by the appearance of colonies on solid media with kanamycin, as pTarget was not eliminated. We determined the escape frequency for the I-OnuI TSM kill switch to be between 10^−4^and 10^−5^(Fig. 4a), similar to reported frequencies for other endonuclease-based kill switches which initiate killing at a single target site^37^. To understand how the kill switch was being inactivated, we isolated total plasmid DNA from 27 escapees and used Oxford Nanopore sequencing to identify mutations in both pEndo and pTarget. A significant fraction of the escape mutants possessed large insertions in pEndo, many of which created frame-shift mutations resulting in truncated endonucleases (Fig. 4b). Interestingly, 10 of these insertions produced a consistent size increase in pEndo from the original 5.2 kb plasmid to a ∼ 6.7 kb plasmid and all insertions were in the VMA1 coding region (Fig. 4c). We identified the insertions as having similarity with IS*911*, a member of the IS*3* bacterial insertion sequence family^38^. IS*911* is known to be temperature-sensitive^39^, increasing in both transposition frequency and production of IS*911*-associated proteins as temperature decreases, suggesting that many of these insertions likely occurred upon shifting the incubation temperature to 18°C. We also identified a hotspot for a common 12-bp deletion in the VMA1 intein sequence that was found in 13 of the escape mutants (Fig. 4b and Fig. 4c). This deletion may occur more frequently in this region due to a 9-bp palindromic region found on either side of the 12-bp VMA1 deletion (Fig. S4), contributing to genetic instability at this locus^40–43^. Only one escape mutant possessed a mutation in pTarget (a 1-bp deletion in the I-OnuI target site); however the same mutant had several indels in pEndo. Many of these indels were found in homopolymeric regions, resulting in uncertainty as to whether these are genuine mutations or sequencing errors^44, 45^.

**Figure 4.**
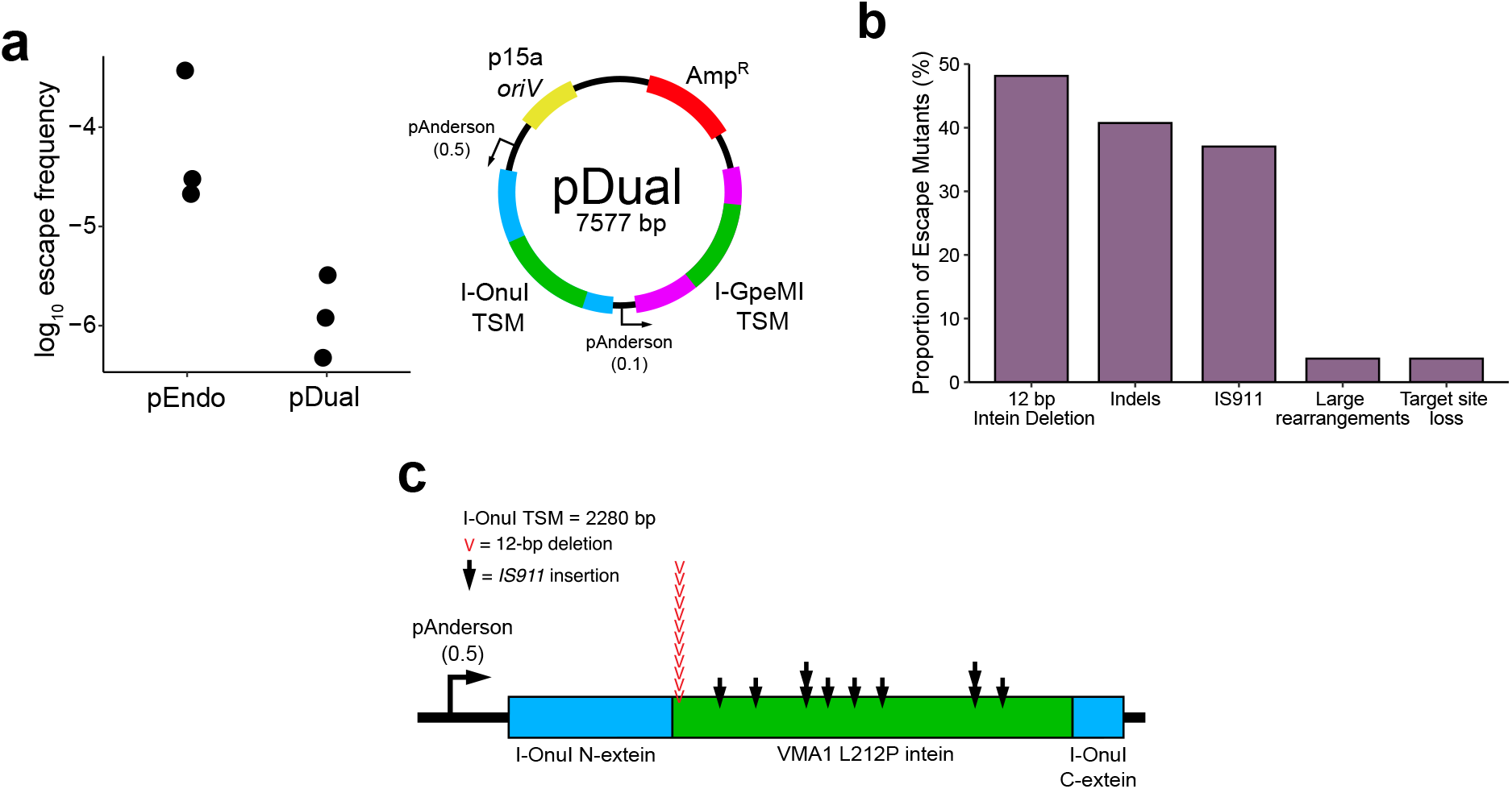
Characterizing escape and inactivation of the I-OnuI TSM. **(A)** Escape frequency from a single- and dual-endonuclease setup. (left) The frequency of TSM inactivation in pEndo and pDual. The escape frequency was determined by the number of colonies grown on solid LB media with kanamycin relative to media without kanamycin, suggesting failure to cleave pTarget. Each data point represents individual replicates (n=3). (right) Schematic of the dual-endonuclease plasmid, pDual. *oriV*, plasmid origin of replication; Amp^R^, ampicillin resistance gene; pAnderson, constitutively active Anderson promoter; TSM, temperature-sensitive meganuclease. Two variations of the Anderson promoter were used, with either low-(0.1) or mid-(0.5) expression relative to the cognate Anderson promoter. Each Anderson promoter controls the expression of a separate TSM open reading frame. **(B)** Sequencing TSM inactivation from escape mutants of pEndo in *E. coli* Nissle 1917. pEndo and pTarget from 27 escape colonies were sequenced using the Oxford Nanopore MinION. Data is shown as a bar plot, with the proportion of each of the listed mutations shown relative to the total number of escape plasmids. **(C)** Mapping the locations of IS*911* insertions and the 12 bp VMA1 intein deletion on the I-OnuI TSM. Black arrows represent IS*911* insertion sites on the I-OnuI TSM open reading frame, and red arrows represent the location of the 12 bp VMA1 intein deletion.

The above analysis revealed that escape mutants arose from indels in the TSM coding region, suggesting that multiplexing TSMs on pEndo could decrease the escape frequency. We constructed a pEndo variant expressing two TSMs by placing the coding region of I-GpeMI with the L212P VMA1 intein inserted at GC2 (I-GpeMI TSM) downstream of the I-OnuI coding region (Fig. 4a). To avoid recombination between genetic elements on pEndo, we used a different variant of the Anderson promoter (BBaJ23100) to express the I-GpeMI TSM and modified the DNA sequence of the L212P VMA1 intein within I-GpeMI, creating a synonymous substitution every third codon. An I-GpeMI target site was also inserted into pTarget. With pDual, we found an *∼* 10-fold decrease in the escape frequency relative to pEndo (Fig. 4a). This result demonstrates that multiplexing orthogonal TSMs to introduce redundancy can reduce the escape frequency.

### The TSM kill switch restricts plasmids to the mouse GI tract

To demonstrate that the I-OnuI and I-GpeMI TSM kill-switches can restrict plasmids to the gut environment, we independently passaged EcN harbouring pEndo or pDual and the appropriate pTarget vector through C57BL/6 mice, collecting fecal samples over three days (Fig. 5a). Both of the EcN strains harbouring active TSMs demonstrated a 10^3^ to 10^5^fold reduction in pTarget retention when fecal resuspensions were incubated at 18°C, observed as a decrease in recovered CFU/mL on kanamycin-containing media (Fig. 5b). No significant reduction in recovered CFU/mL was observed with fecal suspensions from TSM-active strains incubated at 37°C as compared to the E22Q nuclease dead or N454Q intein splicing inactive controls. We found that the escape frequency of each of the pEndo or pDual TSM strains was ~10-fold higher *in vivo* compared to frequencies found for *in vitro* experiments (compare Fig. 5a with Fig. 5b). However, we noted that pDual maintained the 10-fold reduction in escape frequency as compared to pEndo, suggesting that multiplexing orthogonal TSMs is broadly applicable to different environments. Taken together, this data confirms that the TSM kill switch is inactive while in the mouse gut but is activated outside of the mouse gut at 18°C. Furthermore, no side effects were observed in any of the mice, indicating that the circuit appears stable and safe for at least three days in the mouse gut.

**Figure 5.**
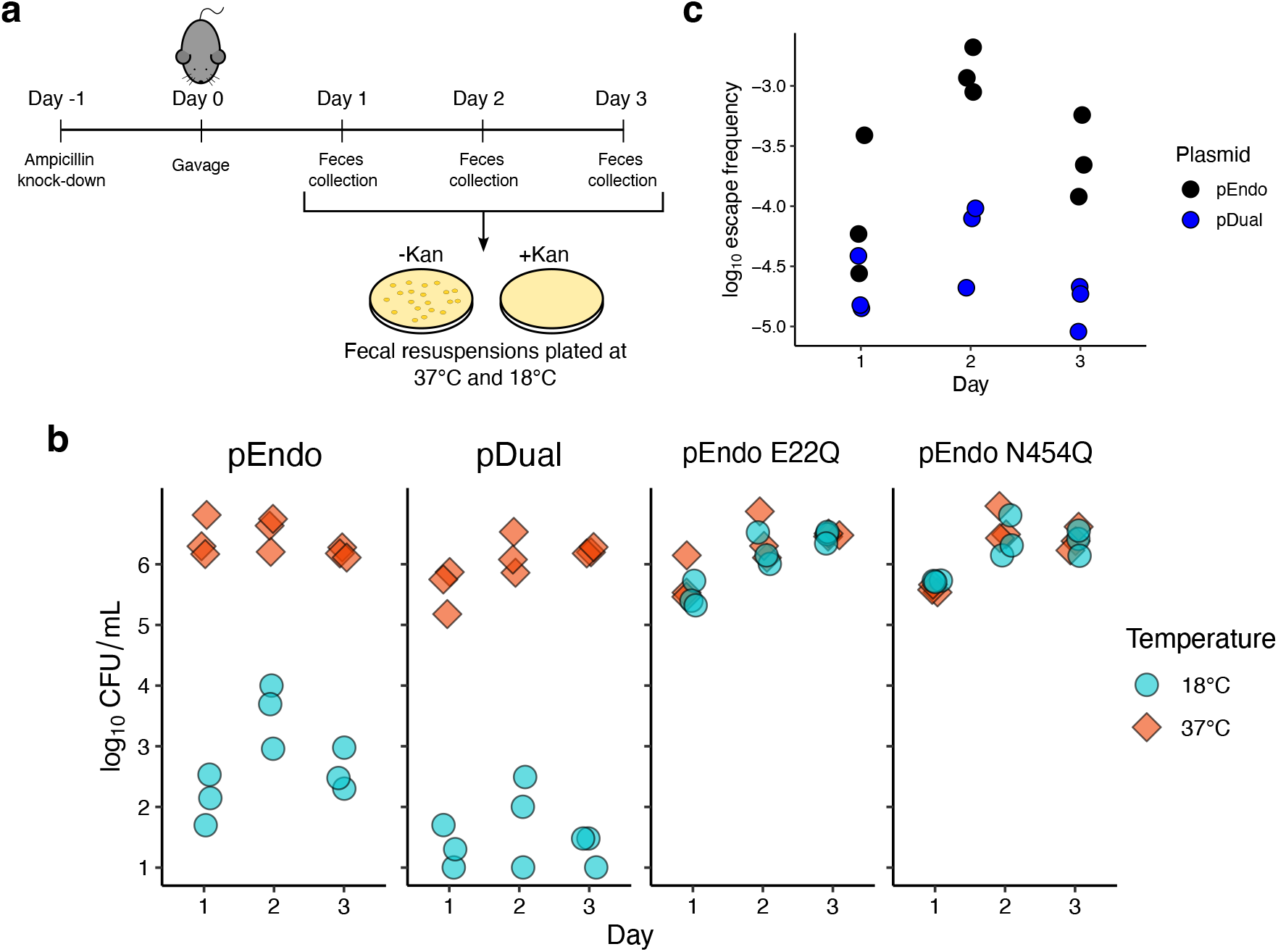
Containment of plasmids with the I-OnuI or I-GpeMI TSMs in the mouse gut. **(A)** Schematic outlining the mouse model experiments. Gut microbiome knockdown was performed with 1 g/L ampicillin and 2.5% sucrose. Mice were gavaged with 10^8^CFUs of pEndo or pDual. Fecal samples were resuspended in PBS (150 *μ*L/mg). **(B)** Depletion of pTarget by pEndo and pDual. Plasmid retention was calculated as the ratio of colonies grown on MacConkey agar with ampicillin and kanamycin, over media with only ampicillin. Each data point represents fecal samples collected from the 3 mice used in each experimental group (n=3). **(C)** Escape frequency of pEndo and pDual in a mouse model. The escape frequency was determined by the ratio of colonies recovered from plates incubated at 18°C relative to colonies recovered at 37°C. Each data point represents individual replicates (n=3).

## Discussion

We created an orthogonal set of thermally-regulated site-specific meganucleases controlled by intein splicing for use as biocontainment kill switches to promote degradation of DNA at low temperatures outside of the mammalian gut, or for other applications where a suitable temperature gradient exists. In contrast to past kill switches that are primarily regulated by transcriptional mechanisms, our system functions at the post-translational level and is controlled by the temperature-sensitive VMA1 L212P intein that, in the context of meganucleases, splices at 20°C or below. The small coding size of meganucleases (under 1-kb) and lack of elaborate transcriptional circuits to control gene expression simplifies both plasmid construction and multiplexing of orthogonal enzymes. Our biocontainment system does not require engineering of the bacterial genome, or require exogenously added signalling molecules to repress or active TSM expression. Because of the high structural conservation between meganuclease family members, other TSMs could be created beyond those tested here by inserting the VMA1 L212P intein into the GC1 and GC2 sites. VMA1 inteins (or other inteins) with different splicing permissive temperatures could be inserted into meganucleases to fine-tune endonuclease activity to a desired temperature depending on the application or organism. This strategy could also be used to create different temperature-controlled site-specific endonucleases, such as the GIY-YIG family endonuclease I-TevI that is permissive to intein insertion^46^, or other toxin systems that promote DNA degradation and bacterial killing.

From a practical viewpoint, the TSM biocontainment system is well suited to experiments that introduce engineered microbes carrying recombinant plasmids into the mammalian GI tract simply because eliminated feces are immediately introduced to a permissive temperature for intein splicing, TSM activation and plasmid degradation. We can envision other applications for TSMs, including the elimination of bacterial strains that are engineered to carry TSM target sites, the curing of plasmids from bacteria, yeast or other species that can grow at a wide range of temperatures, or industrial applications where plasmid-containing cultures can be shifted to low temperatures to induce plasmid destruction. Lastly, layering of TSMs with functionally distinct kill switches could achieve a level of degradation of plasmid or genomic DNA to satisfy biocontainment concerns in therapeutic applications.

## Methods

### Bacterial strains

*E. coli* EPI300 (F^*′*^*λ*^−^*mcrA* ∆(*mrr-hsdRMS-mcrBC*) *ϕ* 80d*lacZδ M15* ∆*(lac)X74 recA1 endA1 araD139* ∆(*ara, leu)7697 galU galK rpsL* (Str^*R*^) *nupG trfA dhfr*) (Epicenter) was used for cloning purposes. *E. coli* NEB 5-alpha (*fhuA2*∆*(argF-lacZ)U169 phoA glnV44 ϕ 80*∆*(lacZ)M15 gyrA96 recA1 relA1 endA1 thi-1 hsdR17*) (New England Biolabs) was used for the initial two-plasmid assay. *E. coli* Nissle 1917 (acquired from Dr. Jeremy Burton at Western University) was used in the characterization experiments, escape assays, and mouse experiments. *E. coli* ER2566 (B F^-^, *λ*^−^*fhuA2*, [*lon*], *ompT lacZ::*T7.1 *gal sulA11* ∆(*mcrC-mrr*) *114*::IS*10* R(*mcr-73*::)miniTN*10*(Tet^S^) *endA1* [*dcm*]) (New England Biolabs) was used for protein expression and purification.

### Plasmid construction

A list of primers is provided in Supplementary Table S1. All plasmids were assembled in *E. coli* EPI300 using either Gibson assembly or Golden Gate Assembly^47, 48^. All Gibson assemblies used the NEBuilder HiFi DNA Assembly Kit (New England Biolabs). All Golden Gate assemblies and mutagenesis were performed using BsmBI (New England Biolabs). Plasmids were designed in Benchling, gene fragments were ordered through Telesis Bio and assembled on the BioXp™3200, and primers were ordered and assembled by Integrated DNA Technologies. Wild-type I-OnuI was previously cloned between the NcoI and NotI cut sites on the pProEX-HT-a plasmid (Invitrogen and Life Technologies) and pEndo backbone^49, 50^.

I-OnuI was removed from the original pEndo vector by inverse PCR, using DE-6667 and DE-5792. I-GpeMI and I-PanMI were cloned into the resulting linear plasmid by Gibson assembly. pEndo I-GpeMI was assembled with the linear pEndo amplicon and DE-6369. pEndo I-PanMI was assembled with the linear pEndo amplicon and DE-6377. pEndo I-GpeMI was linearized at GC1 using DE-6291 and DE-6292, and at GC2 using DE-6293 and DE-6294. The VMA1 intein was cloned by Gibson assembly into GC1 of I-GpeMI using DE-6325 and DE-6326, and at GC2 using DE-6327 and DE-6328. pEndo I-PanMI was linearized at GC1 using DE-6307 and DE-6308, and at GC2 using DE-6309 and DE-6310. The VMA1 intein was cloned by Gibson assembly into GC1 of I-PanMI using DE-6341 and DE-6342, and at GC2 using DE-6343 and DE-6344. pEndo I-OnuI was linearized at GC1 using DE-5706 and DE-5707, and at GC2 using DE-5669 and DE-5670. The VMA1 intein was cloned by Gibson assembly into GC1 of I-OnuI using DE-7157 and DE-7158, and at GC2 using DE-5963 and DE-5964. The E22Q and N454Q mutants of the I-OnuI TSM were created using Golden Gate Mutagenesis^51^using DE-6270 and DE-6271 for E22Q, and DE-6958 and DE-6959 for N454Q at GC2. The pBAD promoter and *araC* gene were removed from original iteration of pEndo I-OnuI using DE-5733 and DE-5734. Both primers contained 35-bp overhangs carrying the sequence for the BBaJ23108 Anderson promoter, which was re-circularized by Gibson assembly. pDual was developed from this pEndo variant by opening a site downstream of the I-OnuI TSM ORF using DE-6963 and DE-6964. The I-GpeMI ORF was cloned out of pEndo I-GpeMI using DE-6961 and DE-6962, excluding the BBaJ23108 Anderson promoter. The I-GpeMI ORF was cloned into pEndo with the BBaJ23100 Anderson promoter using DE-6965, DE-6966, and DE-6960.

pTarget was created from a preexisting toxic plasmid (pTox), as previously described^49, 52^. The *ccdB* DNA gyrase toxin and associated *lac* operon regulatory components were cloned out of pTox using DE-5737 and DE-5738. Both primers carried 20-bp overhangs that allowed it to be re-circularized using Gibson assembly. I-GpeMI and I-PanMI target sites were introduced by cloning out the I-OnuI target site using DE-5830 and DE-5831, and new target sites for I-GpeMI and I-PanMI were inserted using DE-6354 and DE-6359, respectively. pTarget-Dual was developed by linearizing pTarget-Onu using DE-5833 and DE-5834, and introducing an I-GpeMI target site by Gibson assembly using DE-6734.

### In vitro cleavage assays

Wild-type I-OnuI was expressed on pProEX-HT-a harbored in *E. coli* ER2566, cloned between the NcoI/NotI restriction sites. Cultures were grown to mid-log (OD_600_ ∼ 0.5) and and protein expression was induced with 1 mM isopropyl-*β* -D-thiogalactopyranoside (IPTG, BioBasic) overnight at 16°C. Cells were pelleted at 6000 x *g* for 15 minutes, followed by resuspension in binding buffer (50 mM Tris·HCl pH 8.0, 500 mM NaCl, 1 mM imidazole, 10% glycerol) and lysed via sonication. The cell lysate was centrifuged at 16,000 x *g*. The clarified lysate was loaded onto a 1 mL HisTrap-HP column (Cytiva) and washed with 10 mL binding buffer and 10 mL wash buffer (50 mM Tris ·HCl pH 8.0, 500 mM NaCl, 35 mM imidazole, 10% glycerol). I-OnuI was eluted off the column into 5 mL of elution buffer (50 mM Tris· HCl pH 8.0, 500 mM NaCl, 100 mM imidazole, 10% glycerol) and dialyzed against 1 L dialysis buffer (50 mM Tris ·HCl pH 8.0, 250 mM NaCl, 30 mM imidazole, 10% glycerol).

Time-point cleavage assays to determine I-OnuI cleavage activity at different temperatures was performed in reaction buffer (50 mM Tris ·HCl pH 8.0, 100 mM NaCl, 10 mM MgCl_2_, 1 mM dithiothreitol (DTT)), 400 nM purified I-OnuI, and 10 nM supercoiled target plasmid. Reactions were performed in parallel at 37°C, 30°C, 25°C, 22°C, and 18°C. Reactions were stopped at 10 time points using stop solution (100 mM ethylenediaminetetraacetic acid (EDTA), 160 *μ*g Proteinase K (New England Biolabs), dissolved in Dulbecco’s phosphate-buffered saline (D-PBS)). Cleavage products were visualized on a 1% agarose gel and stained with ethidium bromide. The gel was imaged using the ChemiDoc Imaging System (Bio-Rad) and the proportion of nicked, linear, and supercoiled bands were quantified using the Bio-Rad Image Lab software.

### Bacterial two-plasmid assay

We used a modified version of a two-plasmid assay as previously described^52–55^. 50 ng of each pEndo variant was transformed into 50 *μ*L of *E. coli* NEB5*α* carrying pTarget with the appropriate cognate target sites. Cells were allowed to recover in 1 mL 2xYT media (16 g/L tryptone, 10 g/L yeast extract, 5 g/L NaCl) at 37°C and 225 rpm for 1 hour. Following the initial recovery, cells were added to 1 mL of 2X induction media (2xYT, 200 *μ*g/mL carbenicillin, 0.04% arabinose). The media was then split, and 1 mL was induced at 37°C for 1.5 hours and the other 1 mL at 18°C for 2.5 hours. After the outgrowth period, dilutions were prepared and spot plated or spread on M9 minimal media agar plates (1x M9 salts, 0.8% wt/vol tryptone, 1% vol/vol glycerol, 1 mM MgSO_4_, 1 mM CaCl_2_, 0.2% wt/vol thiamine, and 0.02% (wt/vol) L-glucose). Plates were incubated overnight at 37°C or 18°C for 5 days. Colonies were counted manually, and plasmid retention was calculated as the ratio of colonies grown on selective M9 media (carbenicillin (100 *μ*g/mL), kanamycin (50 *μ*g/mL)), to non-selective M9 media (kanamycin (50 *μ*g/mL)). A catalytically inactive I-OnuI mutant (E22Q) and a splicing-deficient VMA1 intein mutant (N454Q) were used as negative controls.

### Time-course cleavage assay

*E. coli* Nissle 1917 (EcN) carrying pEndo with the I-OnuI TSM, E22Q endonuclease knockout, or N454Q intein splicing knockout were grown overnight at 37°C under selection. Overnight cultures were diluted 1:100 into selective LB media, and incubation continued at 18°C for 12 hours. Aliquots of the culture were taken every hour, and plated on non-selective LB agar (carbenicillin (100 *μ*g/mL)) or selective LB (carbenicillin (100 *μ*g/mL), kanamycin (50 *μ*g/mL)). Plates were incubated at either 37°C or 18°C, and colonies were counted manually. Plasmid retention was calculated as previously described.

### Quantitative PCR

Experiments were initiated as previously described for the time-course cleavage assay. After 2 hours of incubation at 18°C, 10 mL of culture was pelleted and resuspended in 500 *μ*L 1X phosphate-buffered saline (PBS). At 12 hours this process was repeated, pelleting 500 *μ*L of culture instead. Samples were boil-lysed at 95°C for 10 minutes, then stored at -20°C. Quantitative real-time PCR was performed using SYBR Select Master Mix (Applied Biosystems), amplifying a 150 bp region of the kanamycin resistance gene on pTarget using DE-7269 and DE-7270, and a 150 bp region of the *CspA* gene in *E. coli* Nissle 1917 using DE-7271 and DE-7272. Three biological and five technical replicates were performed for each sample. Data was plotted as the ratio of pTarget counts to *E. coli* Nissle 1917 chromosome counts. Serial dilutions of purified pTarget and boil-lysed genomic DNA were used as standards.

### Escape mutant analyses

Escape mutants were obtained by passaging *E. coli* Nissle 1917 carrying pEndo with the I-OnuI TSM, E22Q endonuclease knockout, or N454Q intein splicing knockout at 37°C overnight. All cells also carried the appropriate pTarget plasmids. The following day, the culture was diluted 1:100 into fresh LB and incubated at 18°C for another night. The culture was subsequently serially diluted and plated on selective-(100 *μ*g/mL carbenicillin, 50 *μ*g/mL kanamycin) and non-selective (100 *μ*g/mL carbenicillin) LB plates. The escape frequency was calculated as the ratio between the CFU/mL from selective LB incubated at 18°C to the number of colonies from non-selective media. Escape mutant colonies were picked from selective LB plates incubated at 18°C, selecting for failed kill-switches. 30 escape colonies were picked and grown overnight, and plasmids were extracted using the Monarch Plasmid Miniprep Kit (New England Biolabs). Both pEndo and pTarget were transformed into *E. coli* EPI300 to achieve higher plasmid yield after re-isolation. Plasmids were fully-sequenced using the Nanopore MinION (Oxford Nanopore Technologies) by Flow Genomics.

### Mouse experiments

All mouse model experiments used C57BL/6 female mice with three mice per cage. Drinking water and feed were provided *ad libitum*. One day prior to gavage, drinking water was supplemented with ampicillin (1 g/L) and 2.5% sucrose to facilitate knockdown of the gut microbiome. pEndo, pDual, the E22Q endonuclease knockout, and the N454Q intein splicing knockout harbored in *E. coli* Nissle 1917 were grown overnight in selective LB at 37°C. On the day of gavage, the overnight cultures were diluted 1:50 into fresh LB media, and grown to mid-log (OD_600_ ∼ 0.5). The cells were pelleted and resuspended with 1X PBS, concentrating the cells to 10^8^CFUs/100 *μ*L. Each mouse was gavaged with 100 *μ*L of the appropriate sample. For three days following gavage, mice fecal pellets were collected daily and resuspended in PBS (150 *μ*L/mg) by vortexing and mechanical agitation. Samples were serially diluted and plated on selective-(100 *μ*g/mL carbenicillin, 50 *μ*g/mL kanamycin) and non-selective (100 *μ*g/mL carbenicillin) MacConkey agar. Colonies were counted manually to determine the CFU/mL for each sample and both temperatures. The escape frequency was calculated as the reduction between the CFU/mL for pEndo and pDual from plates grown at 18°C compared to 37°C.

## Funding

Supported by a Project Grant (PJT-159708) to D.R.E. from the Canadian Institutes of Health Research.

## Acknowledgements

We thank Jeremy Burton for providing the *E. coli* Nissle 1917 strain.

## Notes

### Competing Interest Statement

The authors have declared no competing interest.

