## Supplementary Data for "Intein-based thermoregulated meganucleases for biocontainment of genetic material"

**This document includes:**
Figs. S1-S5

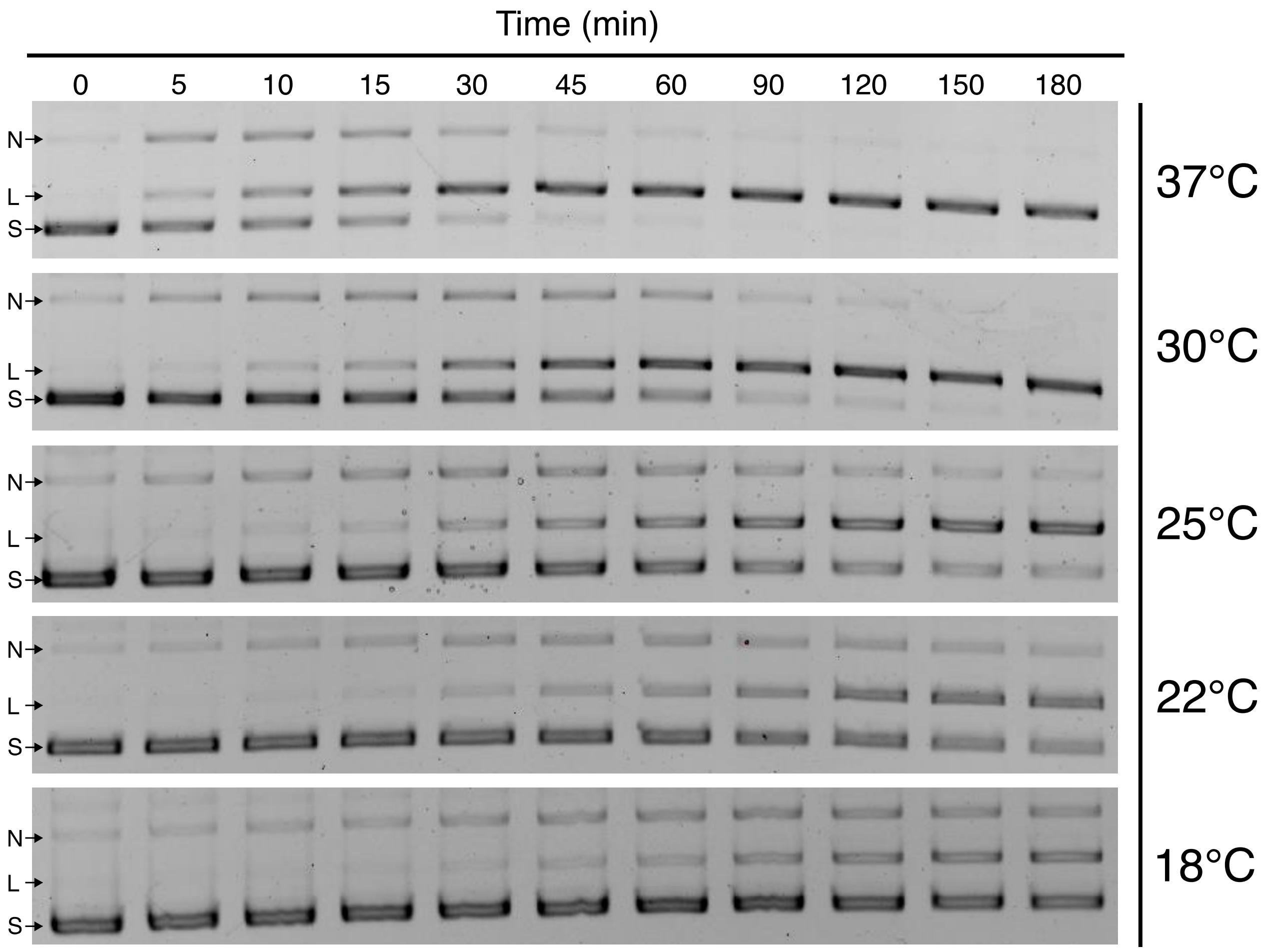


Figure S1. *in vitro* cleavage activity of wild-type I-OnuI against supercoiled plasmid incubated different temperatures. Labeled are the nicked (N), linear (L), and supercoiled (S) bands on each gel. Images cropped for publication.

| | |
|---|---|
| I-OnuI | TTTCCACTTATTCAACCTTTTA |
| I-AniI | CTGAGGAGGTTTCTCTGTAAAG |
| I-CkaMI | AAGTACCCTTTAAACCTAATTA |
| I-GpeMI | TTTCCGCTTATTCAACCCTTTA |
| I-GpiI | TTTTCCTGTATATGACTTAAAT |
| I-GzeI | GCCCCTCATAACCCGTATCAAG |
| I-GzeII | ATGGGTACCATATTGGTACAAA |
| I-PanMI | GCTCCTCATAATCCTTATCAAG |
| I-SmaMI | TATCCTCCATTATCAGGTGTAC |
| I-SscMI | AGGTACCCTTTAAACCTATTAA |

Figure S2. Cognate DNA substrates for 10 LAGLIDADG homing endonucleases. Highlighted in red are the “critical four” residues, for which nucleotide substitutions are least tolerated.

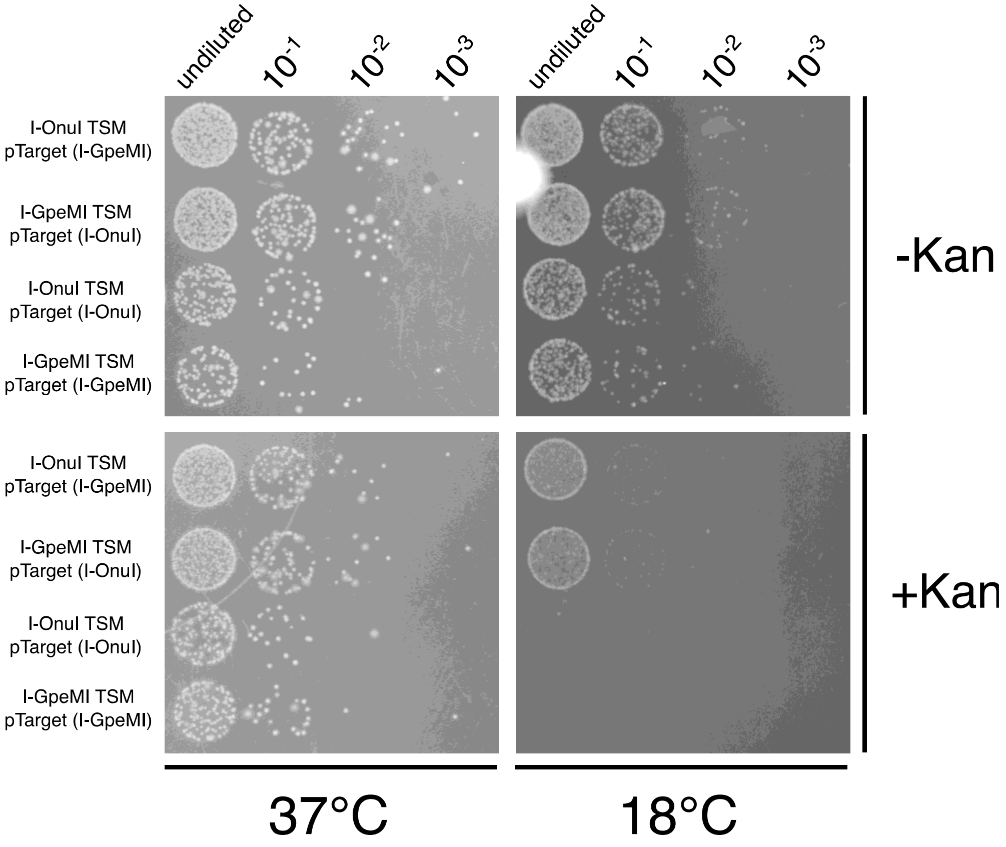


Figure S3. Cleavage preference orthogonality of I-OnuI and I-GpeMI. Cleavage is indicated by the failure to grow on kanamycin-supplemented media, resulting from the loss of pTarget.

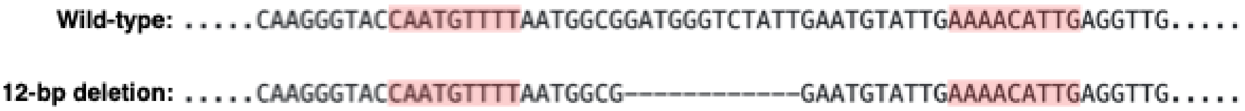


Figure S4. 12-bp deletion product resulting in frequent I-OnuI TSM inactivation. Highlighted in red are the palindromic inverted repeat regions thought to contribute to the observed frequency of the deletion.

>VMA1
CFAKGTNVLMADGSIECIENIEVGNKVMGKDGRPREVIKLPRGRETMYSVVQKSQHRA
HKSDSSREVPELLKFTCNATHELVVRTPRSVRRLSRTIKGVEYFEVITFEMGQKKAPDG
RIVELVKEVSKSYPISEGPERANELVESYRKASNKAYFEWTIEARDLSLLGSHVRKATYQ
TYAPILYENDHFFDYMQKSKFHLTIEGPKVLAYLLGLWIGDGLSDRATFSVDSRDTSLME
RVTEYAEKLNLCAEYKDRKEPQVAKTVNLYSKVVRGNGIRNNLNTENPLWDAIVGLGFL
KDGVKNIPSFLSTDNIGTRETFLAGLIDSDGYVTDEHGIKATIKTIHTSVRDGLVSLARSL
GLVVSVNAEPAKVDMNGTKHKISYAIYMSGGDVLLNVLSKCAGSKKFRPAPAAAFAREC
RGFYFELQELKEDDYYGITLSDDSDHQFLLANQVVVHN

>WT I-OnuI
MGSAYMSRRESINPWILTGFADAEGSFLLRIRNNNKSSVGYSTELGFQITLHNKDKSILE
NIQSTWKVGVIANSGDNAVSLKVTRFEDLKVIIDHFEKYPLITQKLGDYMLFKQAFCVME
NKEHLKINGIKELVRIKAKLNWGLTDELKKAFPEIISKERSLINKNIPNFKWLAGFTSGEG
CFFVNLIKSKSKLGVQVQLVFSITQHIKDKNLMNSLITYLGCGYIKEKNKSEFSWLDFVV
TKFSDINDKIIPVFQENTLIGVKLEDFEDWCKVAKLIEEKKHLTESGLDEIKKIKLNMNKG
RVF*

>I-OnuI/VMA1 (GC1)
MGSAYMSRRESINQWILTGFADAEGSFLLRIRNNNKSSVGYSTELGFQITLHNKDKSILE
NIQSTWKVGVIANSGDNAVSLKVTRFEDLKVIIDHFEKYPLITQKLGDYMLFKQAFCVME
NKEHLKINGIKELVRIKAKLNWGLTDELKKAFPEIISKERSLINKNIPNFKWLAGFTSGEG
CFAKGTNVLMADGSIECIENIEVGNKVMGKDGRPREVIKLPRGRETMYSVVQKSQHRA
HKSDSSREVPELLKFTCNATHELVVRTPRSVRRLSRTIKGVEYFEVITFEMGQKKAPDG
RIVELVKEVSKSYPISEGPERANELVESYRKASNKAYFEWTIEARDLSLLGSHVRKATYQ
TYAPILYENDHFFDYMQKSKFHLTIEGPKVLAYLLGLWIGDGLSDRATFSVDSRDTSLME
RVTEYAEKLNLCAEYKDRKEPQVAKTVNLYSKVVRGNGIRNNLNTENPLWDAIVGLGFL
KDGVKNIPSFLSTDNIGTRETFLAGLIDSDGYVTDEHGIKATIKTIHTSVRDGLVSLARSL
GLVVSVNAEPAKVDMNGTKHKISYAIYMSGGDVLLNVLSKCAGSKKFRPAPAAAFAREC
RGFYFELQELKEDDYYGITLSDDSDHQFLLANQVVVHNCFFVNLIKSKSKLGVQVQLVF
SITQHIKDKNLMNSLITYLGCGYIKEKNKSEFSWLDFVVTKFSDINDKIIPVFQENTLIGVK
LEDFEDWCKVAKLIEEKKHLTESGLDEIKKIKLNMNKGRVF*

>I-OnuI/VMA1 (GC2)
MGSAYMSRRESINQWILTGFADAEGSFLLRIRNNNKSSVGYSTELGFQITLHNKDKSILE
NIQSTWKVGVIANSGDNAVSLKVTRFEDLKVIIDHFEKYPLITQKLGDYMLFKQAFCVME
NKEHLKINGIKELVRIKAKLNWGLTDELKKAFPEIISKERSLINKNIPNFKWLAGFTSGEG
CFFVNLIKSKSKLGVQVQLVFSITQHIKDKNLMNSLITYLGCFAKGTNVLMADGSIECIEN
IEVGNKVMGKDGRPREVIKLPRGRETMYSVVQKSQHRAHKSDSSREVPELLKFTCNAT
HELVVRTPRSVRRLSRTIKGVEYFEVITFEMGQKKAPDGRIVELVKEVSKSYPISEGPER
ANELVESYRKASNKAYFEWTIEARDLSLLGSHVRKATYQTYAPILYENDHFFDYMQKSK
FHLTIEGPKVLAYLLGLWIGDGLSDRATFSVDSRDTSLMERVTEYAEKLNLCAEYKDRK
EPQVAKTVNLYSKVVRGNGIRNNLNTENPLWDAIVGLGFLKDGVKNIPSFLSTDNIGTR
ETFLAGLIDSDGYVTDEHGIKATIKTIHTSVRDGLVSLARSLGLVVSVNAEPAKVDMNGT
KHKISYAIYMSGGDVLLNVLSKCAGSKKFRPAPAAAFARECRGFYFELQELKEDDYYGI
TLSDDSDHQFLLANQVVVHNCGYIKEKNKSEFSWLDFVVTKFSDINDKIIPVFQENTLIG
VKLEDFEDWCKVAKLIEEKKHLTESGLDEIKKIKLNMNKGRVF*

>I-OnuI(E22Q)/VMA1 (GC2)
MGSAYMSRRESINQWILTGFADAQGSFLLRIRNNNKSSVGYSTELGFQITLHNKDKSIL
ENIQSTWKVGVIANSGDNAVSLKVTRFEDLKVIIDHFEKYPLITQKLGDYMLFKQAFCVM
ENKEHLKINGIKELVRIKAKLNWGLTDELKKAFPEIISKERSLINKNIPNFKWLAGFTSGE
GCFFVNLIKSKSKLGVQVQLVFSITQHIKDKNLMNSLITYLGCFAKGTNVLMADGSIECIE
NIEVGNKVMGKDGRPREVIKLPRGRETMYSVVQKSQHRAHKSDSSREVPELLKFTCN
ATHELVVRTPRSVRRLSRTIKGVEYFEVITFEMGQKKAPDGRIVELVKEVSKSYPISEGP
ERANELVESYRKASNKAYFEWTIEARDLSLLGSHVRKATYQTYAPILYENDHFFDYMQK
SKFHLTIEGPKVLAYLLGLWIGDGLSDRATFSVDSRDTSLMERVTEYAEKLNLCAEYKD
RKEPQVAKTVNLYSKVVRGNGIRNNLNTENPLWDAIVGLGFLKDGVKNIPSFLSTDNIG
TRETFLAGLIDSDGYVTDEHGIKATIKTIHTSVRDGLVSLARSLGLVVSVNAEPAKVDMN
GTKHKISYAIYMSGGDVLLNVLSKCAGSKKFRPAPAAAFARECRGFYFELQELKEDDYY
GITLSDDSDHQFLLANQVVVHNCGYIKEKNKSEFSWLDFVVTKFSDINDKIIPVFQENTLI
GVKLEDFEDWCKVAKLIEEKKHLTESGLDEIKKIKLNMNKGRVF*

>I-OnuI/VMA1(N454Q) (GC2)
MGSAYMSRRESINQWILTGFADAEGSFLLRIRNNNKSSVGYSTELGFQITLHNKDKSILE
NIQSTWKVGVIANSGDNAVSLKVTRFEDLKVIIDHFEKYPLITQKLGDYMLFKQAFCVME
NKEHLKINGIKELVRIKAKLNWGLTDELKKAFPEIISKERSLINKNIPNFKWLAGFTSGEG
CFFVNLIKSKSKLGVQVQLVFSITQHIKDKNLMNSLITYLGCFAKGTNVLMADGSIECIEN
IEVGNKVMGKDGRPREVIKLPRGRETMYSVVQKSQHRAHKSDSSREVPELLKFTCNAT
HELVVRTPRSVRRLSRTIKGVEYFEVITFEMGQKKAPDGRIVELVKEVSKSYPISEGPER
ANELVESYRKASNKAYFEWTIEARDLSLLGSHVRKATYQTYAPILYENDHFFDYMQKSK
FHLTIEGPKVLAYLLGLWIGDGLSDRATFSVDSRDTSLMERVTEYAEKLNLCAEYKDRK
EPQVAKTVNLYSKVVRGNGIRNNLNTENPLWDAIVGLGFLKDGVKNIPSFLSTDNIGTR
ETFLAGLIDSDGYVTDEHGIKATIKTIHTSVRDGLVSLARSLGLVVSVNAEPAKVDMNGT
KHKISYAIYMSGGDVLLNVLSKCAGSKKFRPAPAAAFARECRGFYFELQELKEDDYYGI
TLSDDSDHQFLLANQVVVHQCGYIKEKNKSEFSWLDFVVTKFSDINDKIIPVFQENTLIG
VKLEDFEDWCKVAKLIEEKKHLTESGLDEIKKIKLNMNKGRVF*

>WT I-GpeMI
MGPTRNESINPWVLTGFADAEGSFILRIRNNNKSSAGYSTELGFQITLHKKDISILENIQS
TWKVGVIANSGDNAVSLKVTRFEDLRVVLNHFEKYPLITQKLGDYLLFKQAFSVMENKE
HLKIEGIKRLVGIKANLNWGLTDELKEAFVASGGENIFVASGGERSLINKNIPNSGWLAG
FTSGEGCFFVSLIKSKSKLGVQVQLVFSITQHARDRALMDNLVTYLGCGYIKEKKKSEF
SWLEFVVTKFSDIKDKIIPVFQVNNIIGVKLEDFEDWCKVAKLIEEKKHLTESGLEEIRNIK
LNMNKGRVL*

>I-GpeMI/VMA1 (GC1)
MGPTRNESINPWVLTGFADAEGSFILRIRNNNKSSAGYSTELGFQITLHKKDISILENIQS
TWKVGVIANSGDNAVSLKVTRFEDLRVVLNHFEKYPLITQKLGDYLLFKQAFSVMENKE
HLKIEGIKRLVGIKANLNWGLTDELKEAFVASGGENIFVASGGERSLINKNIPNSGWLAG
FTSGEGCFAKGTNVLMADGSIECIENIEVGNKVMGKDGRPREVIKLPRGRETMYSVVQ
KSQHRAHKSDSSREVPELLKFTCNATHELVVRTPRSVRRLSRTIKGVEYFEVITFEMGQ
KKAPDGRIVELVKEVSKSYPISEGPERANELVESYRKASNKAYFEWTIEARDLSLLGSH
VRKATYQTYAPILYENDHFFDYMQKSKFHLTIEGPKVLAYLLGLWIGDGLSDRATFSVDS

RDTSLMERVTEYAEKLNLCAEYKDRKEPQVAKTVNLYSKVVRGNGIRNNLNTENPLWD
AIVGLGFLKDGVKNIPSFLSTDNIGTRETFLAGLIDSDGYVTDEHGIKATIKTIHTSVRDGL
VSLARSLGLVVSVNAEPAKVDMNGTKHKISYAIYMSGGDVLLNVLSKCAGSKKFRPAPA
AAFARECRGFYFELQELKEDDYYGITLSDDSDHQFLLANQVVVHNCFFVSLIKSKSKLG
VQVQLVFSITQHARDRALMDNLVTYLGCGYIKEKKKSEFSWLEFVVTKFSDIKDKIIPVF
QVNNIIGVKLEDFEDWCKVAKLIEEKKHLTESGLEEIRNIKLNMNKGRVL*

>I-GpeMI/VMA1 (GC2)
MGPTRNESINPWVLTGFADAEGSFILRIRNNNKSSAGYSTELGFQITLHKKDISILENIQS
TWKVGVIANSGDNAVSLKVTRFEDLRVVLNHFEKYPLITQKLGDYLLFKQAFSVMENKE
HLKIEGIKRLVGIKANLNWGLTDELKEAFVASGGENIFVASGGERSLINKNIPNSGWLAG
FTSGEGCFFVSLIKSKSKLGVQVQLVFSITQHARDRALMDNLVTYLGCFAKGTNVLMAD
GSIECIENIEVGNKVMGKDGRPREVIKLPRGRETMYSVVQKSQHRAHKSDSSREVPEL
LKFTCNATHELVVRTPRSVRRLSRTIKGVEYFEVITFEMGQKKAPDGRIVELVKEVSKSY
PISEGPERANELVESYRKASNKAYFEWTIEARDLSLLGSHVRKATYQTYAPILYENDHFF
DYMQKSKFHLTIEGPKVLAYLLGLWIGDGLSDRATFSVDSRDTSLMERVTEYAEKLNLC
AEYKDRKEPQVAKTVNLYSKVVRGNGIRNNLNTENPLWDAIVGLGFLKDGVKNIPSFLS
TDNIGTRETFLAGLIDSDGYVTDEHGIKATIKTIHTSVRDGLVSLARSLGLVVSVNAEPAK
VDMNGTKHKISYAIYMSGGDVLLNVLSKCAGSKKFRPAPAAAFARECRGFYFELQELK
EDDYYGITLSDDSDHQFLLANQVVVHNCGYIKEKKKSEFSWLEFVVTKFSDIKDKIIPVF
QVNNIIGVKLEDFEDWCKVAKLIEEKKHLTESGLEEIRNIKLNMNKGRVL*

>WT I-PanMI
MGFKRNFSTLESKLNPSYISGFVDGEGSFMLTIIKDNKYKLGWRVVCRFVISLHKKDLSL
LNKIKEFFDVGNVFLMTKDSAQYRVESLKGLDLIINHFDKYPLITKKQADYKLFKMAHNLI
KNKSHLTKEGLLELVAIKAVINNGLNNDLSIAFPGINTILRPDTSLPQILNPFWLSGFVDAE
GCFSVVVFKSKTSKLGEAVKLSFILTQSNRDEYLIKSLIEYLGCGNTSLDPRGTIDFKVTN
FSSIKDIIVPFFIKYPLKGNKNLDFTDFCEVVRLMENKSHLTKEGLDQIKKIRNRMNTNRK
*

>I-PanMI/VMA1 (GC1)
MGFKRNFSTLESKLNPSYISGFVDGEGSFMLTIIKDNKYKLGWRVVCRFVISLHKKDLSL
LNKIKEFFDVGNVFLMTKDSAQYRVESLKGLDLIINHFDKYPLITKKQADYKLFKMAHNLI
KNKSHLTKEGLLELVAIKAVINNGLNNDLSIAFPGINTILRPDTSLPQILNPFWLSGFVDAE
GCFAKGTNVLMADGSIECIENIEVGNKVMGKDGRPREVIKLPRGRETMYSVVQKSQHR
AHKSDSSREVPELLKFTCNATHELVVRTPRSVRRLSRTIKGVEYFEVITFEMGQKKAPD
GRIVELVKEVSKSYPISEGPERANELVESYRKASNKAYFEWTIEARDLSLLGSHVRKATY
QTYAPILYENDHFFDYMQKSKFHLTIEGPKVLAYLLGLWIGDGLSDRATFSVDSRDTSLM
ERVTEYAEKLNLCAEYKDRKEPQVAKTVNLYSKVVRGNGIRNNLNTENPLWDAIVGLGF
LKDGVKNIPSFLSTDNIGTRETFLAGLIDSDGYVTDEHGIKATIKTIHTSVRDGLVSLARSL
GLVVSVNAEPAKVDMNGTKHKISYAIYMSGGDVLLNVLSKCAGSKKFRPAPAAAFAREC
RGFYFELQELKEDDYYGITLSDDSDHQFLLANQVVVHNCFSVVVFKSKTSKLGEAVKLS
FILTQSNRDEYLIKSLIEYLGCGNTSLDPRGTIDFKVTNFSSIKDIIVPFFIKYPLKGNKNLD
FTDFCEVVRLMENKSHLTKEGLDQIKKIRNRMNTNRK*

>I-PanMI/VMA1 (GC2)
MGFKRNFSTLESKLNPSYISGFVDGEGSFMLTIIKDNKYKLGWRVVCRFVISLHKKDLSL

LNKIKEFFDVGNVFLMTKDSAQYRVESLKGLDLIINHFDKYPLITKKQADYKLFKMAHNLI
KNKSHLTKEGLLELVAIKAVINNGLNNDLSIAFPGINTILRPDTSLPQILNPFWLSGFVDAE
GCFSVVVFKSKTSKLGEAVKLSFILTQSNRDEYLIKSLIEYLGCFAKGTNVLMADGSIECI
ENIEVGNKVMGKDGRPREVIKLPRGRETMYSVVQKSQHRAHKSDSSREVPELLKFTC
NATHELVVRTPRSVRRLSRTIKGVEYFEVITFEMGQKKAPDGRIVELVKEVSKSYPISEG
PERANELVESYRKASNKAYFEWTIEARDLSLLGSHVRKATYQTYAPILYENDHFFDYMQ
KSKFHLTIEGPKVLAYLLGLWIGDGLSDRATFSVDSRDTSLMERVTEYAEKLNLCAEYK
DRKEPQVAKTVNLYSKVVRGNGIRNNLNTENPLWDAIVGLGFLKDGVKNIPSFLSTDNI
GTRETFLAGLIDSDGYVTDEHGIKATIKTIHTSVRDGLVSLARSLGLVVSVNAEPAKVDM
NGTKHKISYAIYMSGGDVLLNVLSKCAGSKKFRPAPAAAFARECRGFYFELQELKEDDY
YGITLSDDSDHQFLLANQVVVHNCGNTSLDPRGTIDFKVTNFSSIKDIIVPFFIKYPLKGN
KNLDFTDFCEVVRLMENKSHLTKEGLDQIKKIRNRMNTNRK*

Figure S5. Amino acid sequences of homing endonucleases and TSMs.
